# novelBGC: An interactive dual-score framework for biosynthetic gene cluster novelty assessment and candidate prioritisation

**DOI:** 10.64898/2026.06.15.732227

**Authors:** Gyanesh Shukla, Balavardhan Merugu, Gaurav Sharma

## Abstract

Genome mining now yields tens of thousands of putative biosynthetic gene clusters (BGCs) per project, yet, separating genuinely novel candidates from rediscoveries of known compounds remains the rate-limiting step before experimental validation. Single-axis prioritisation tools, antiSMASH similarity, BiG-FAM GCF distance, and self-resistance-enzyme (SRE) filters such as ARTS, each surface a different facet of evidence, yet their isolated use systematically over-ranks rediscovery-prone BGCs and overlooks genuinely orphan clusters. We present novelBGC, a web-hosted framework that converts these disparate outputs into two deliberately non-inverse continuous metrics per BGC, a Novelty (N) and a Reference Similarity (RS) score which together define a 2D decision plane that resolves rediscoveries, divergent family members, contig-edge artefacts, and uncharted chemistry with interactive visualisations, with all component weights user-tuneable at submission. Retrospective validation across three independent experimental datasets demonstrates the utility of the framework for candidate prioritization. Within the first 186-BGC SRE-guided cloning study, every confirmed bioactive product fell within the low-to-mid N band whereas 55 high-N (N ≥ 0.50) BGCs were never selected. Moreover, in the other two studies, it correctly prioritised the fully orphan lariocidin BGC of *Paenibacillus sp*. M2 and the divergent within-family indanopyrrole-A idp BGC of *Streptomyces sp*. CNX-425. Together, these case studies demonstrate that the joint (N, RS) space facilitates prioritization decisions that are difficult to achieve using any single criterion alone. from identical input data. novelBGC requires no command-line expertise, no local tool installation, and no manual integration of intermediate output formats, addressing a well-documented accessibility barrier for wet-laboratory researchers engaging with genome-mining workflows. novelBGC is freely available at https://project.iith.ac.in/sharmaglab/novelbgc/.

## Introduction

Natural products (NPs) constitute the chemical foundation of many essential therapeutics, including antibiotics, antifungals, immunosuppressants, and anticancer agents, the majority of which are derived from or inspired by microbial biosynthetic pathways (Newman & Cragg, 2020). Historically, NP discovery relied on bioactivity-guided screening of microbial extracts, a process both time-consuming and prone to the repeated re-identification of known compounds. The advent of next-generation sequencing technologies owing to the dramatic reduction in sequencing costs transformed this paradigm by enabling genome-scale interrogation of biosynthetic potential (Katz & Baltz, 2016; Zhang et al., 2017). Genome mining the systematic bioinformatic identification of biosynthetic gene clusters (BGCs) encoding secondary metabolite pathways allows researchers to map the full chemical potential of a microorganism without requiring laboratory expression of the pathway (Baltz, 2021). Pioneering tools such as antiSMASH (Blin et al., 2021) and PRISM (Skinnider et al., 2020), which employ Hidden Markov Models (HMMs) and profile HMMs to detect conserved biosynthetic core enzymes and signature gene architectures, revealed that most sequenced microbial genomes harbor far more BGCs than could be identified and anticipated from classical screening alone, fundamentally changing the scale at which NP discovery could be pursued (Bauman et al., 2021; Lee et al., 2020; Salim et al., 2025).

The success of these identification tools, combined with the exponential growth of publicly available microbial genome sequences, rapidly generated a new and pressing challenge. While the JGI IMG-ABC (Hadjithomas et al., 2015) catalogued over one million automatically predicted BGCs by 2017, modern databases have scaled dramatically. Today, the latest antiSMASH database (Blin et al., 2026) contains 497,429 BGCs, BiG-FAM (Kautsar, Blin, et al., 2021) features a clustered network of roughly 1.2 million BGCs, and the recently launched Secondary Metabolism Collaboratory (SMC) database by JGI holds a staggering 13.1 million BGC regions (Udwary et al., 2025). Despite these massive numbers of computationally predicted clusters, the vast majority remain “orphan” clusters with no known chemical product (Shukla & Sharma, 2024). In stark contrast, the latest version of the gold-standard repository for experimentally characterized clusters, MIBiG 4.0 (Zdouc et al., 2025), contains only 3,059 entries. Consequently fundamental bottleneck in NP discovery has shifted, depicting that the limiting step is no longer identifying BGCs, but selecting the most promising candidates for the costly and labour-intensive downstream experimental work required to isolate and the utilitcharacterize a novel compound (Scherlach & Hertweck, 2021). Compounding this challenge is the persistent problem of rediscovery, the repeated re-identification of BGCs encoding well-known scaffolds such as non-ribosomal peptides and polyketides which consumes experimental resources without advancing the chemical frontier (Albarano et al., 2020).

To straighten this entangled thread of prioritisation challenges, several strategies have been developed to address the prioritisation challenge. Target-directed genome mining, grounded in the resistance hypothesis, exploits the observation that antibiotic-producing organisms must encode self-protection mechanisms resistance genes, immunity determinants, or duplicated copies of the antibiotic target that frequently co-localize within the producing BGC (Almabruk et al., 2018). The Antibiotic-Resistant Target Seeker (ARTS) (Mungan et al., 2020) automates this principle by integrating BGC prediction with detection of resistance determinants and essential housekeeping gene duplications, thereby enriching for BGCs with antibiotic biosynthetic potential. In parallel, comparative genomics approaches have enabled large-scale dereplication by grouping homologous BGCs into Gene Cluster Families (GCFs). Tools such as BiG-SCAPE (Navarro-Muñoz et al., 2020) and BiG-SLiCE (Kautsar, Van Der Hooft, et al., 2021) organize BGCs from public databases into GCF networks BiG-SLiCE, for instance, clustered over 1.2 million BGCs across the BiG-FAM (Kautsar, Blin, et al., 2021) reference database enabling rapid determination of whether a query BGC belongs to a known family or represents an uncharacterized lineage. Pipeline frameworks such as BGCFlow (Weber & Palsson, 2024) further orchestrate these individual tools into end-to-end analytical workflows. Despite their powerful analysis in the backend, these resources share three limiting characteristics that constrain their broader adoption: they require command-line expertise and dedicated bioinformatics infrastructure to deploy; their outputs are distributed across incompatible formats and separate result files that must be manually integrated; and, critically, none provides a standardised, transparent scoring scheme that quantifies BGC novelty as a continuous, evidence-weighted metric integrating multiple evidence streams simultaneously.

Here we present novelBGC, a web server that addresses this gap by providing researchers with a unified, interactive platform for BGC novelty assessment that requires no bioinformatics expertise to operate. novelBGC automatically integrates outputs from antiSMASH (Blin et al., 2021), BiG-SLiCE, and CARD-RGI (Alcock et al., 2020) to compute two complementary continuous scores for each predicted BGC. The Reference Similarity score (RS ∈ [0,1]) quantifies the degree of evidence that a BGC belongs to a well-characterized biosynthetic family, integrating direct sequence homology to MIBiG-curated reference clusters (Terlouw et al., 2023), positional distance to known GCFs in BiG-FAM, and the presence of antimicrobial resistance genes as a functional annotation signal. The Novelty score (N ∈ [0,1]) captures divergence from all known biosynthetic chemistry, incorporating multiplicative penalties when circumstantial evidence such as GCF affiliation with a characterized family or co-localized resistance genes suggests a functional link to known compound classes despite low direct sequence similarity. Crucially, RS and N are not mathematical inverses of one another: a BGC may score moderately on both metrics when it is a peripheral member of a GCF whose best-characterized representative shares only partial similarity. This two-dimensional scoring space is more informative than any single ranking axis and allows researchers to tailor candidate selection to their specific objectives, whether seeking new structural variants of known scaffolds or pursuing entirely unprecedented biosynthetic chemistry. All scoring weights are user-adjustable, and results are presented through an interactive visualisation interface enabling intuitive filtering, comparison, and export of prioritized candidates. Here we describe the novelBGC server architecture, scoring algorithm, and web interface, case studies, and demonstrate its application to the prioritisation of biosynthetic diversity across a set of microbial genomes.

## Materials and Methods

### Data sources and preprocessing

novelBGC accepts raw genomic sequences in FASTA format as primary input through a web-based upload interface. To ensure standardised file handling, the platform restricts uploads to files with .fna or .fasta extensions. Both complete genome assemblies and draft assemblies are supported without restriction. Upon submission, users assign descriptive nomenclature to each genome following a structured ‘Genus_species_strain’ naming convention, which facilitates systematic organization and tracking. Uploaded genomes are queued for processing through the novelBGC computational pipeline using a Redis-backed Celery distributed task queue, which enables asynchronous job execution and maintains web interface responsiveness during computationally intensive analyses.

The pipeline initiates with genome quality assessment using QUAST (Quality Assessment Tool for Genome Assemblies) (Gurevich et al., 2013), which evaluates assembly metrics including contig statistics, N50 values, total assembly length, and GC content. This quality control step ensures that downstream biosynthetic gene cluster prediction is performed being aware of BGC calls arising from fragmented or low-quality assemblies. Genomes after quality assessment proceed to BGC identification using antiSMASH (Antibiotics and Secondary Metabolite Analysis Shell), the current gold standard for automated biosynthetic gene cluster detection. The antiSMASH module identifies BGC regions based on the presence of signature biosynthetic enzymes and characteristic gene architectures associated with known secondary metabolite classes. The resulting antiSMASH outputs undergo additional processing through custom scripts developed for novelBGC, which systematically extract protein sequences encoded within the boundaries of each predicted BGC region. These extracted protein sequences are submitted to CARD-RGI (Comprehensive Antibiotic Resistance Database Resistance Gene Identifier) for antimicrobial resistance gene detection, and the antiSMASH output folders are provided as input for BiG-SLiCE analysis of individual BGCs against the BiG-FAM reference database.

### Comparative BGC annotation and contextualization using genomics pipeline

The novelBGC platform implements a multi-tiered comparative analysis pipeline to comprehensively assess the novelty and functional potential of predicted biosynthetic gene clusters. At the sequence level, the platform employs the KnownClusterBlast module, utilizing the underlying ClusterBlast algorithm, to perform robust, homology-based comparisons against the MIBiG database (Terlouw et al., 2023), which is the premier repository of experimentally characterized BGCs. This analysis quantifies sequence similarity at the cluster level, generating percentage identity scores that facilitate the inference of potential chemical scaffolds when high-similarity matches are detected. To contextualize these predictions within the broader landscape of global biosynthetic diversity, novelBGC leverages BiG-SLiCE (Kautsar, Van Der Hooft, et al., 2021) to calculate the distance between each query BGC and the centroids of approximately 1.2 million BGCs organized into GCFs within the BiG-FAM database (Kautsar, Blin, et al., 2021). Small centroid distances indicate shared biosynthetic mechanisms with established families, whereas large distances suggest unprecedented gene architectures and novel chemistry. To enhance the interpretability of these assignments, custom scripts interrogate the closest matching GCF for the presence of MIBiG-validated reference clusters inside the GCFs (Alas et al., 2024); GCFs containing known representatives provide higher confidence for functional predictions, while those lacking characterized members represent prime, unexplored targets for discovery. Complementing these cluster-level comparisons, the antimicrobial resistance gene screening through CARD-RGI represents a critical component of the prioritisation framework, as the presence of resistance genes or duplicated copies of antibiotic target genes within BGC boundaries often indicates clusters with antibiotic biosynthetic potential. This analysis leverages the well-established resistance hypothesis, which posits that organisms producing antimicrobial compounds must encode self-protection mechanisms to avoid suicide during compound biosynthesis. Ultimately, all these orthogonal evidence streams are unified within a customisable, weighted aggregation model, allowing researchers to dynamically adjust the prioritisation parameters to favour either the discovery of entirely novel pathways or the identification of new variants within known compound classes.

### Dual-scoring (RS-N score) framework for BGC novelty assessment

To systematically quantify and rank biosynthetic gene clusters for experimental characterisation, novelBGC implements a dual-scoring framework that computes two complementary continuous metrics for each predicted BGC. The Reference Similarity score (RS) provides a confidence estimate that a given BGC belongs to a well-characterized biosynthetic family with known chemical products, while the Novelty score (*N*) assesses the likelihood that the BGC encodes a unique chemical scaffold divergent from previously described natural products. Critically, these two scores are not mathematical inverses of one another: a BGC may score moderately on both metrics when, for example, it is a member of a GCF that lacks experimentally characterized representatives. This two-dimensional representation provides a richer prioritisation space than a single novelty axis, enabling researchers to select candidates based on their specific research objectives, whether seeking new variants of known scaffolds or pursuing entirely unprecedented biosynthetic chemistry. The scoring model integrates five evidence components derived from the comparative analysis pipeline, all normalised to a [0,1] scale or encoded as binary indicators to ensure commensurability prior to aggregation. Direct sequence homology is captured via MIBiG Sequence Similarity (*S*_*mibig*_), which equals to the percentage similarity score normalised to [0,1] if a significant MIBiG hit is detected within the BGC boundaries, and is set to 0 otherwise. To quantify the query BGC’s position within the global landscape of biosynthetic diversity, GCF distance components are calculated using a core membership threshold of *T* = 900. Based on the BiG-SLiCE distance *d*, two complementary piecewise-linear functions are derived: the proximity function δ(*d*), which decays from 1 (*d* ≤ *T*) to 0 (*d* > 2*T*) to support the Reference Similarity score, and the divergence function ν(*d*), which inversely increases from 0 to 1 to support the Novelty score. Contextual evidence is integrated through two binary indicators and a structural penalty. Reference presence in GCF (*I*_*ref*_) equals 1 when the closest matching GCF contains experimentally characterized MIBiG clusters (Alas et al., 2024), while AMR Gene Presence (*I*_*amr*_) equals 1 when CARD-RGI detects antimicrobial resistance genes within the BGC. Both indicators elevate similarity confidence while acting as multiplicative penalties on novelty, reflecting the principle that BGCs retaining hallmarks of known biosynthetic classes are less likely to encode entirely unprecedented chemistry. Finally, a Contig Edge Penalty (*P*_*edge*_) addresses the reduced reliability of incomplete BGCs by applying a 20% confidence discount (*P*_*edge*_ = 0.8) to both scores for assemblies located at contig termini, whereas internal BGCs remain unpenalized (*P*_*edge*_ = 1.0). The final scores are computed by aggregating these normalised components. The Reference Similarity score (*RS*) is calculated as a weighted mean of the similarity-oriented metrics, scaled by the edge penalty factor:

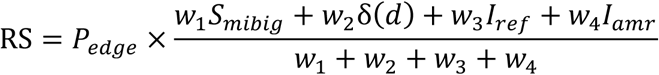

The default weights (*w*_1_ = 5.0, *w*_2_ = 3.0, *w*_3_ = 1.5, *w*_4_ = 1.0) prioritize direct sequence homology and core GCF membership as the strongest indicators of functional annotation. Conversely, the Novelty calculation first establishes a base score (N_base_) reflecting sequence and positional divergence:

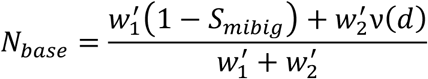

using default weights of 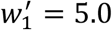 and 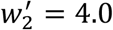. The final Novelty score (N) is then derived by applying the contextual penalties (α = 0.25, β = 0.15) and the structural edge discount:

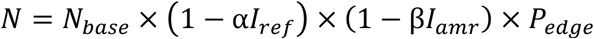

All parameters and penalty coefficients are user-adjustable at the time of submission, allowing the model to be tailored to specific research priorities. Both final scores are clamped to the [0,1] interval and reported to two decimal places.

### Web server implementation and computational infrastructure

The novelBGC web server is implemented as a Flask-based application that orchestrates a comprehensive, multi-tool computational pipeline. At the core of the backend infrastructure, Nextflow workflow management coordinates the optimised, parallel execution of primary analysis components, including QUAST, antiSMASH, BiG-SLiCE, and CARD-RGI. To ensure seamless interoperability among these heterogeneous tools, a suite of custom Python scripts performs essential data transformations, parsing intermediate outputs and precisely structuring inputs for downstream modules. This modular design not only streamlines the current workflow but also readily accommodates the future integration of additional analytical tools. Asynchronous job scheduling and task management are handled by a Redis-backed Celery distributed task queue operating within an Ubuntu server environment, ensuring the user interface remains responsive during intensive background computations. Finally, the application is deployed via a Gunicorn WSGI HTTP server behind an Nginx reverse proxy, providing a robust configuration that efficiently manages concurrent client requests, secure load distribution, and overall system scalability.

### Interactive visualisation and exploratory analytics

The novelBGC results interface employs interactive data visualisation techniques implemented through D3.js and Plotly libraries to facilitate intuitive exploration of complex biosynthetic gene cluster analysis results. The visualisation suite encompasses diverse chart types including pie charts, donut charts, treemaps, interactive scatter plots, and bar plots, all rendered with a carefully selected color palette designed to enhance data interpretation and user experience.

### User interface and analytical workflow

The novelBGC web server provides a streamlined, accessible platform organized into submission, tracking, and interactive results exploration (Figure 1). Users submit genomic assemblies in FASTA format alongside organism name. An advanced options panel is available to customize the scoring weights of the novelty assessment criteria based on specific research priorities. Upon submission, a unique Job ID is generated to track execution and retrieve results, which are retained on the server for 30 days. For users wishing to evaluate the platform before uploading their own data, a pre-computed sample genome is also provided.

**Figure 1.**
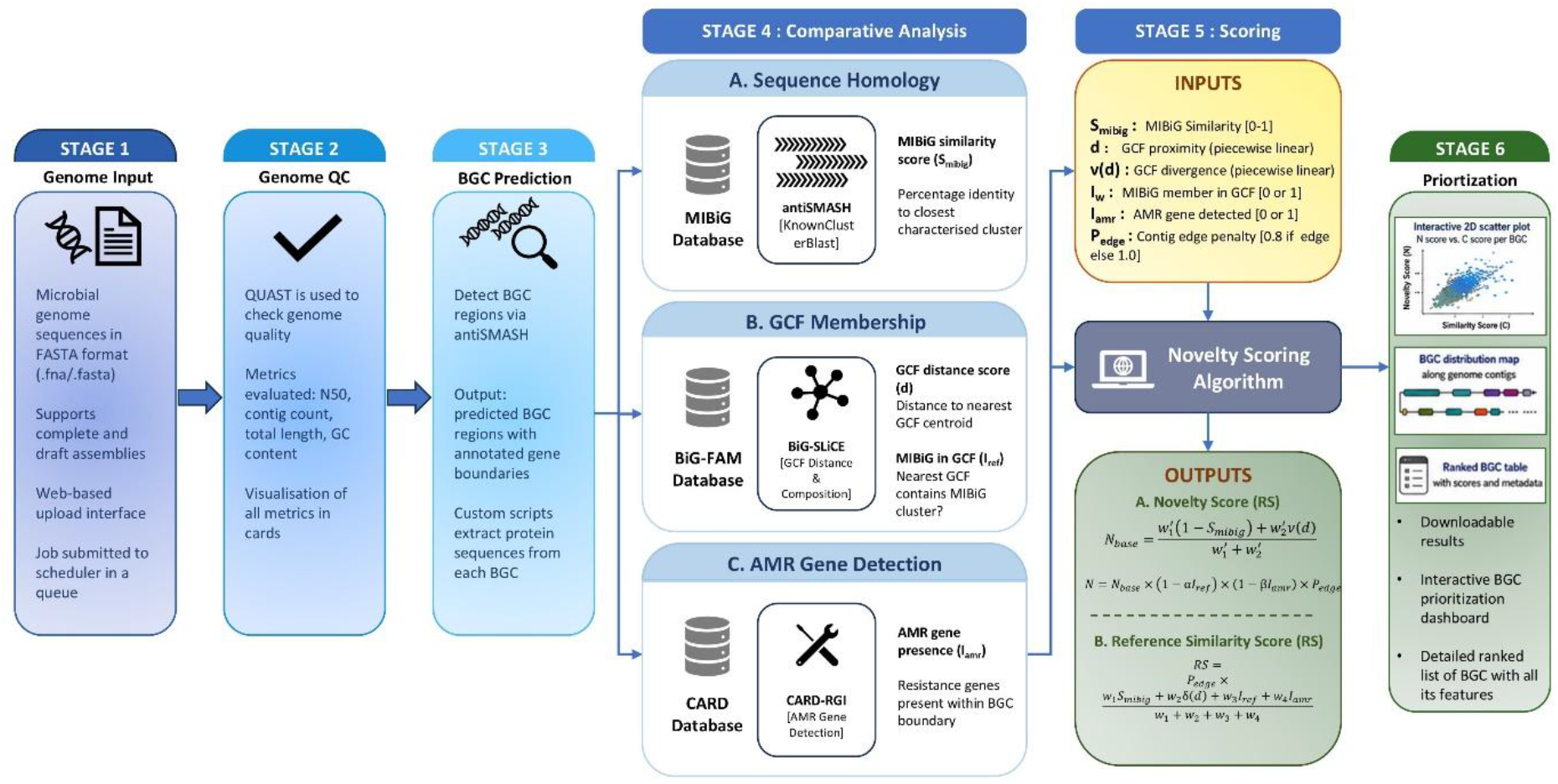
Overview of the novelBGC workflow for biosynthetic gene cluster (BGC) novelty scoring and prioritisation. The pipeline begins with genome assemblies provided in FASTA format (Stage 1), followed by genome quality assessment using QUAST (Stage 2). High-quality genomes proceed to BGC prediction using antiSMASH (Stage 3). In Stage 4, comparative analyses are performed through three parallel modules: (i) sequence homology analysis against the MIBiG, (ii) gene cluster family (GCF) membership analysis, and (iii) antimicrobial resistance (AMR) gene detection against the CARD database. These features are integrated in Stage 5 by the novelty scoring algorithm to calculate two complementary metrics: Reference Similarity (RS) and Novelty Score (N). Finally, Stage 6 provides interactive visualisation for prioritisation of candidate BGCs

To guide researchers through progressively detailed levels of analysis, the interactive results interface is divided into functional modules. The first module provides a high-level overview of genome statistics, detailing assembly quality metrics such as contiguity and genome size to contextualize the overall biosynthetic potential (Figure 2). Alongside this, summary cards quickly distill key information, reporting BGC class distributions, MIBiG similarity counts, edge-contig status, and the presence of antimicrobial resistance (AMR) genes. The second module focuses on spatial distribution, mapping the predicted BGC regions across the assembly contigs. This visualisation is paired with a comprehensive, interactive data table that reports all BGC metadata alongside their calculated Novelty (N) and Reference Similarity (RS) scores (Figure 3). Within this table, users can inspect individual BGC architectures directly via an embedded antiSMASH viewer, which includes options to export specific regions in SVG or GenBank formats.

**Figure 2.**
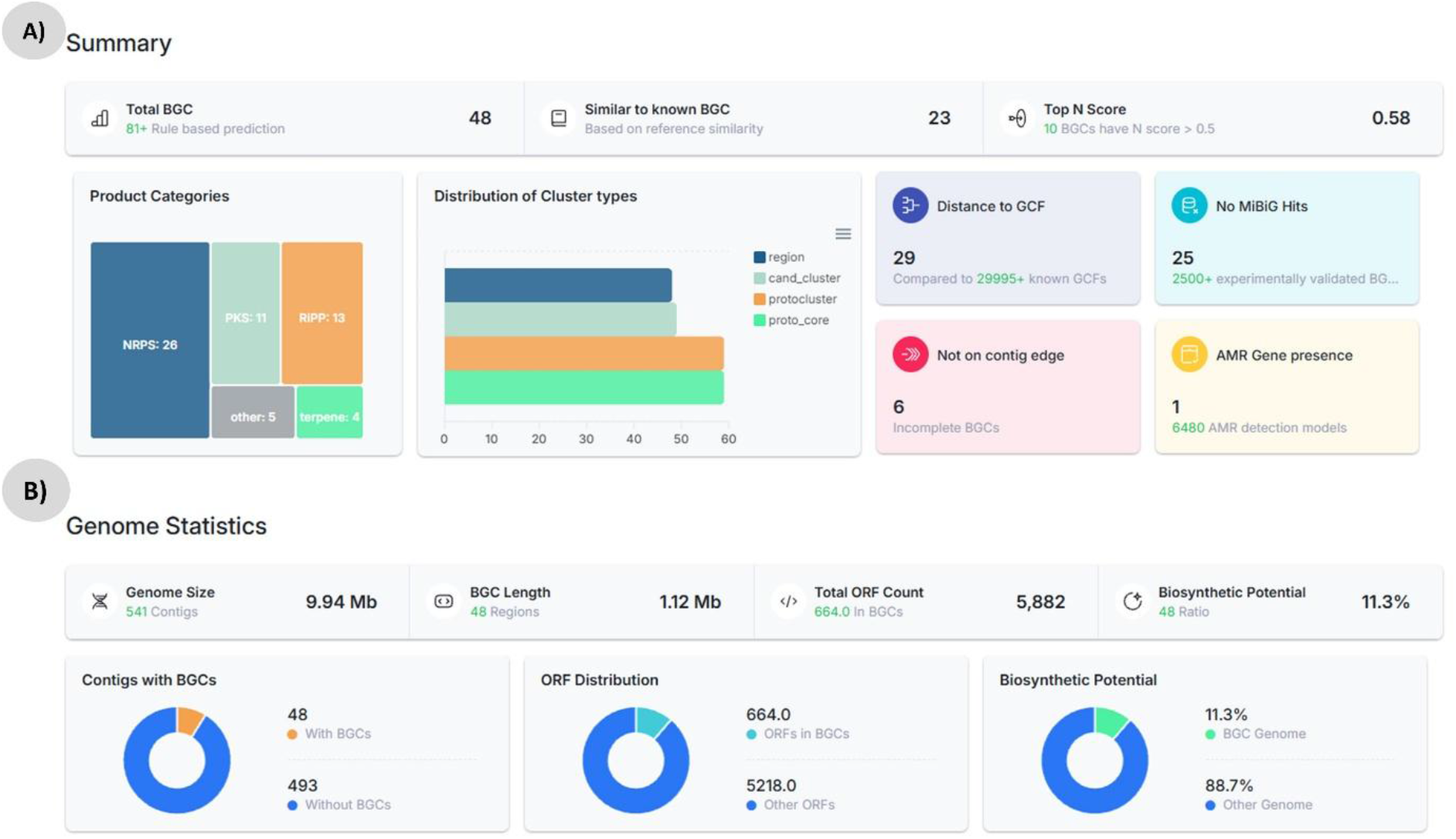
Dashboard style visualisation of result page, divided into 4 distinct sections. All cards have tooltips explaining the meaning of each metric (triggered on hovering over card icon) A) The summary section shows the overall statistics of the processed genome. B) Genome statistics sections show metrics analysed from outputs of QUAST and antiSMASH.

**Figure 3.**
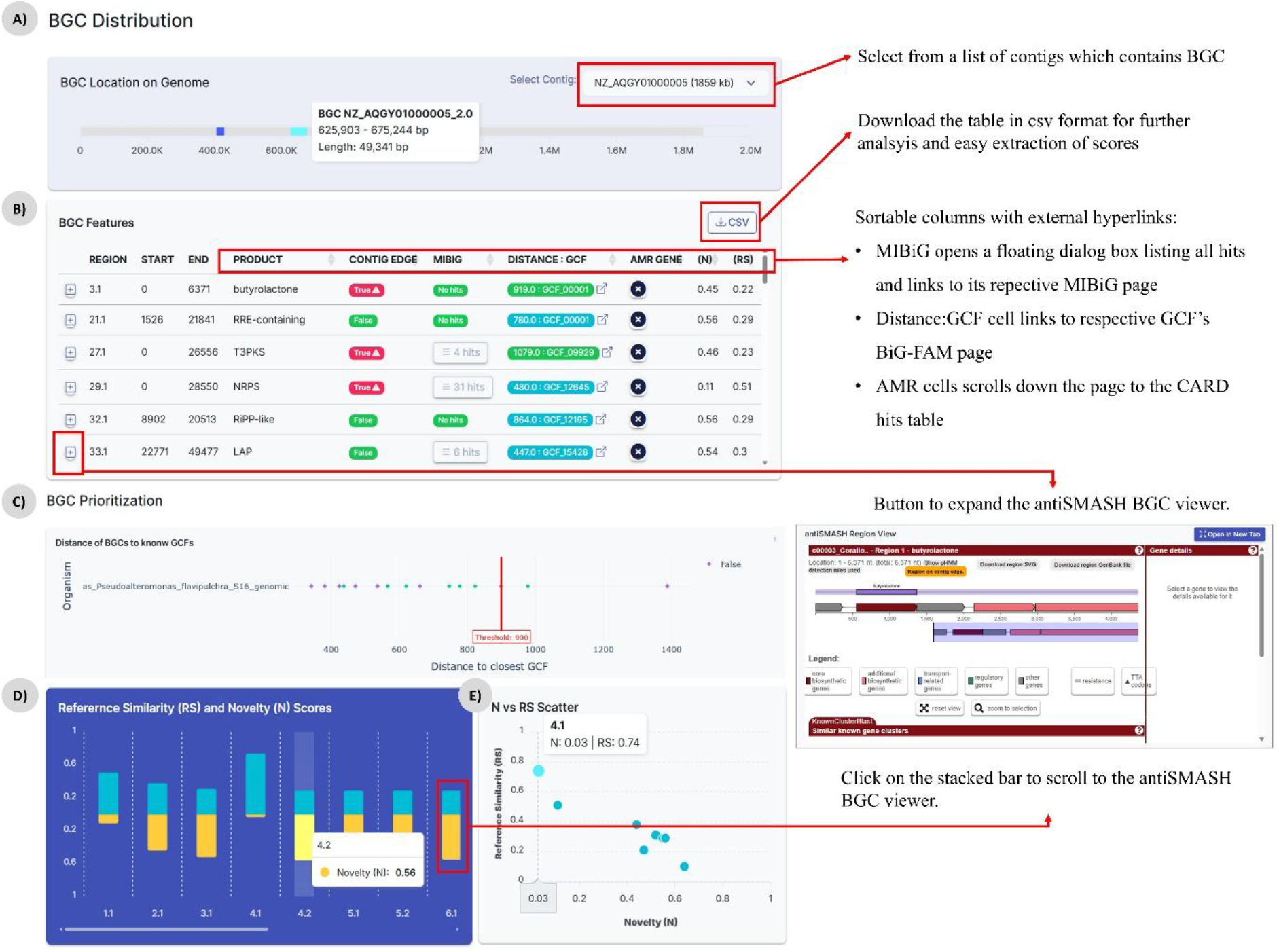
BGC distribution and prioritisation modules. A) BGC location map showing the contigs in which BGCs reside. B) A comprehensive BGC features table giving in depth information about, product, BGC boundary, similarity to MIBiG cluster, distance to closest GCF, resistance gene presence, N-RS scores. C) Strip chart of distance of BGCs to closest GCF, hovering over the jitter shows more metadata info about the BGC and GCF it corresponds to. D) Stacked bar plot of Novelty score (blue) and Reference Similarity score (yellow) of each BGC. E) Scatter plot of N-RS score of all BGCs of the processed genome.

The core analytical outputs are presented in the subsequent module, which visualizes BGC prioritisation to guide candidate selection. This section features a strip chart plotting BGCs against their closest GCF distances to highlight a distance threshold of 900, a stacked bar chart comparing N and RS scores on one axis, and a two-dimensional scatter plot positioning candidates within the novelty-similarity landscape (Figure 3). If AMR determinants are detected, an integrated CARD-RGI table details the relevant resistance mechanisms and associated antibiotics. Finally, the platform concludes with a data export section, enabling the seamless download of complete, publication-ready analysis archives, including QUAST, antiSMASH, and CARD-RGI outputs, as well as execution logs to ensure transparency and facilitate offline integration.

### Retrospective validation across diverse discovery paradigms

To evaluate the predictive accuracy and biological utility of the novelBGC dual-scoring framework, we performed retrospective analyses across three distinct experimental scenarios: bulk dereplication of known compounds, untargeted discovery of unprecedented chemical scaffolds, and targeted pattern-based mining. These analyses demonstrate novelBGC’s capacity to synthesize diverse orthogonal data streams into actionable biological insights that extend significantly beyond the simple aggregation of component tool outputs.

### Case study 1: Bulk dereplication and overcoming single-criterion blind spots

High-throughput genome mining campaigns frequently struggle with rediscovery rates and false positives when relying on single-criterion selection. To demonstrate novelBGC’s ability to navigate this challenge, we evaluated the expert-curated FAST-NPS dataset (Yuan et al., 2025), wherein 105 BGCs from 11 *Streptomyces* strains were selected for expression solely based on the presence of self-resistance enzymes (SREs) identified via ARTS (Mungan et al., 2020). While SRE localization is a powerful metric, using it in isolation can lead to rediscovery traps. For example, novelBGC correctly assigned a low Novelty score (N = 0.13) and high Reference Similarity score (RS = 0.65) to a linaridin BGC in *S. bikiniensis* (region 12.3, 77% similarity to cypemycin) that was subsequently experimentally confirmed to produce that known compound. Similarly, three BGCs selected by FAST-NPS via SREs, *S. nojiriensis* region 1.6 (96% similarity to JBIR-126; N = 0.02, RS = 0.74), *S. spororaveus* region 5.23 (96% similarity to JBIR-126; N = 0.02, RS = 0.74), and *S. sp*. NRRL F-5635 region 1.2 (93% similarity to venemycin; N = 0.04, RS = 0.73) were each flagged by novelBGC as near-certain rediscovery risks before any experimental commitment. Furthermore, novelBGC isolated two SRE-triggered false positives, *S. spororaveus* region 5.6 (2-methylisoborneol, 100 % MIBiG; N = 0.00, RS = 0.76) and *S. nojiriensis* region 1.9 (desferrioxamine B, 100 % MIBiG; N = 0.00, RS = 0.76). 2-Methylisoborneol is the ubiquitous geosmin-class earthy-odour volatile of soil Streptomyces, with no reported antibacterial or antitumor activity (Becher et al., 2020), and desferrioxamine B is a near-universal actinomycete hydroxamate siderophore used clinically as the iron-chelator Desferal (Knapp & Giddings, 2026) rather than as an antibiotic. Because neither metabolite engages an intracellular target of the producer, no self-resistance enzyme is biologically required and the resistance-gene context detected by ARTS reflects a coincidental paralog of a housekeeping enzyme rather than a genuine SRE-HKE pairing, unlike the 77-96 % rediscoveries above, which remain within the antibiotic chemotype space the SRE filter was designed to enrich.

To ensure this dereplication capability is robust across diverse, independently mined genomes, we expanded our analysis beyond the FAST-NPS dataset to include the recent marine isolate *Streptomyces sp*. CNX-435 (Sweeney et al., 2024) (the source of indanopyrrole A, discussed below). Within this genome, novelBGC correctly assigned absolute baseline novelty scores (N = 0) and high RS scores to regions encoding ubiquitous, well-characterized metabolites such as ectoine (Region 1.2), alongside other known clusters including E-837 (Region 15.1) and citrulassin D (Region 10.2) (Supplementary Table S2). By reliably flagging these regions, novelBGC provides a robust, population-level dereplication metric that simultaneously prevents wasted experimental effort and clears genomic “noise.”

### Case study 2: Prioritizing unprecedented chemistry: the Lariocidin breakthrough

While dereplication is critical, the primary goal of many untargeted discovery programs is the identification of entirely unprecedented chemical scaffolds within genomic “dark matter.” In standard annotation pipelines, highly divergent BGCs often return zero MIBiG hits, making it difficult for researchers to distinguish between non-functional evolutionary remnants and highly valuable novel clusters. To test novelBGC’s predictive power in this context, we retrospectively applied it to the genome of *Paenibacillus sp*. M2, which was recently reported to produce lariocidin, a novel lassopeptide representing a completely new structural family that targets an immutable ribosomal site (Jangra et al., 2025). Without any prior knowledge of the publication or specific structural biases, novelBGC successfully isolated the exact productive BGC from the genomic background, assigning it the highest Novelty score of 0.61in the entire genome and an exceptionally low RS score of 0.25 (Supplementary Table S3). This specific dual-score profile mathematically mirrored the biological reality of the discovery: the maximal N-score accurately quantified the unprecedented structural divergence of the lariocidin family, while the low RS-score predicted the molecule’s lack of known resistance mechanisms (Table 1, row 6) (Supplementary Table S1).

**Table 1.**
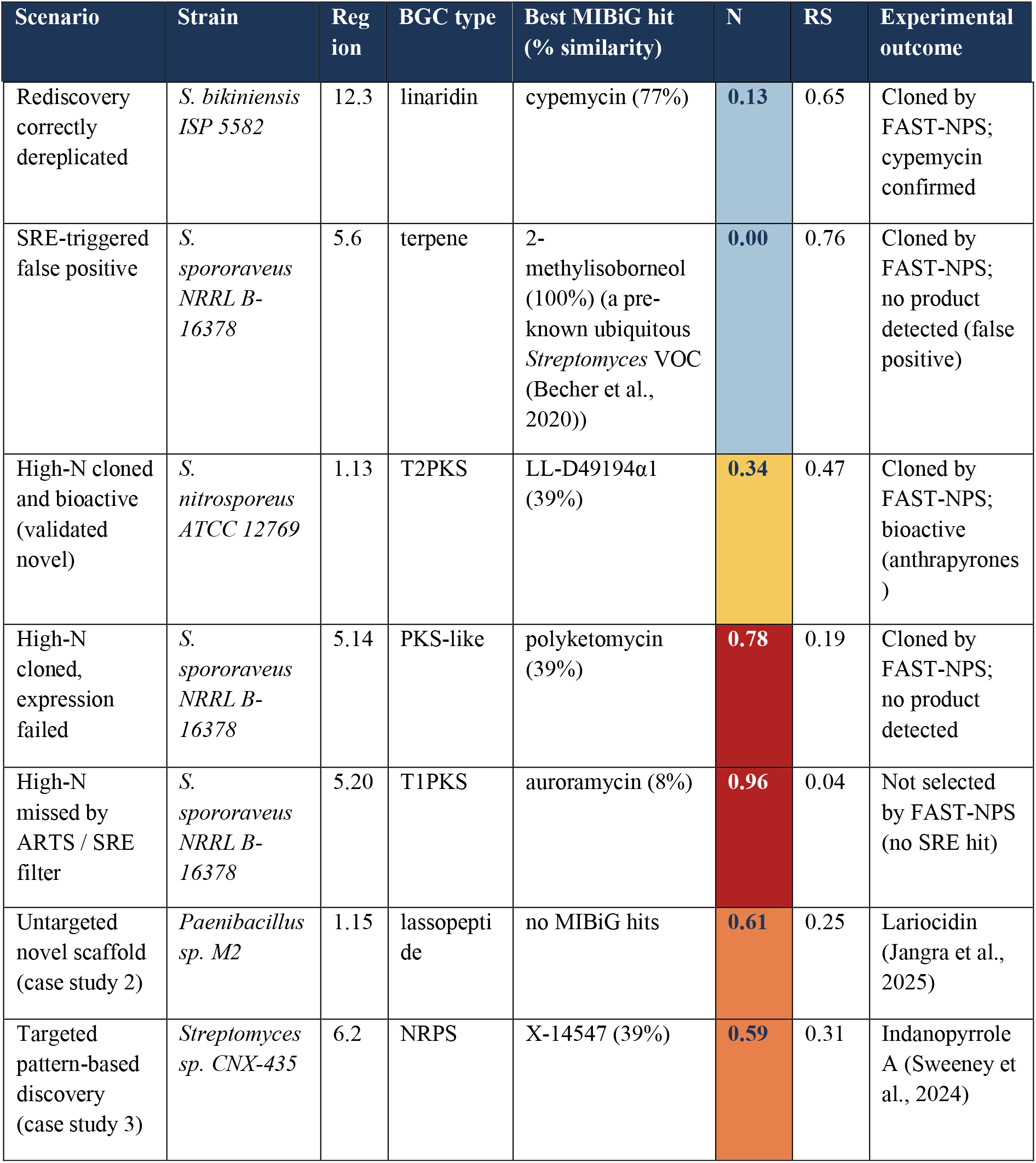
Representative BGCs spanning the seven outcome scenarios encountered across the three retrospective case studies. Each row anchors a numerical claim made in case studies and links it to a single BGC; the full dataset is provided as Supplementary Table S1. N = Novelty score; RS = Reference Similarity score. % sim. = MIBiG cluster similarity reported by KnownClusterBlast. Rows 1-5 are drawn from the FAST-NPS dataset (cf. Figure 4); row 6 from *Paenibacillus sp*. M2 (case study 2); row 7 from *Streptomyces sp*. CNX-435 (case study 3).

### Case study 3: Navigating targeted discovery and algorithmic scope

While maximum N-scores effectively highlight entirely uncharted biosynthetic space, practical genome mining is frequently guided by targeted, pattern-based searches for specific structural motifs rather than genome-wide novelty. We note that novelBGC’s ranking reflects biosynthetic divergence as defined by GCF distance, presence of MIBiG member in closest GCF, AMR gene presence in BGC and MIBiG similarity; it does not encode experimental tractability, structural accessibility, or a research group’s specific scaffold interests. For example, in the recent characterisation of the marine *Streptomyces sp*. CNX-435 (Sweeney et al., 2024), authors successfully discovered the novel antibiotic indanopyrrole A from region 6.2. When analysed by novelBGC, region 6.2 was correctly identified as a high-novelty, low-reference-similarity candidate (N = 0.59, RS = 0.31), successfully marking it as highly divergent, however, it ranked 10th overall by N-score within that specific genome (Supplementary Table S2). The experimental prioritisation of this BGC by the original authors was correctly guided by targeted pattern-based mining for the indanopyrrole scaffold, not by maximising global novelty. In such targeted campaigns, novelBGC serves to computationally showcase beforehand that a biologically selected candidate might possesses sufficient divergence from known databases to justify their experimental efforts, though researchers will have to apply additional domain-specific filters beyond the N-score alone.

## Discussion

The principal advance of novelBGC is the conversion of disparate, evidence-rich bioinformatics outputs into a single, interpretable two-dimensional prioritisation landscape. By computing the Reference Similarity (RS) and Novelty (N) scores as deliberately non-inverse continuous metrics, the framework allows researchers to distinguish four operationally distinct quadrants, confirmed rediscoveries, divergent members of known families, contig-edge artefacts, and entirely uncharted chemistry, within a single visualisation, rather than collapsing this structure onto a single rank. Across the three retrospective case studies presented here, this dual-score representation consistently delivered actionable predictions that other existing single-criterion approaches could not produce from the same genome inputs. The dereplication, untargeted-discovery, and targeted-mining scenarios summarised in Table 1 (Supplementary Table S1) each surfaced a distinct failure mode of single-axis prioritisation; novelBGC successfully addressed all three scenarios examined here by anchoring its predictions to the position of each BGC in the joint (N, RS) space rather than to its rank along either axis alone.

The FAST-NPS analysis illustrates this most concretely (Figure 4). Self-resistance enzyme localisation remains a powerful guide for antibiotic biosynthesis, but in isolation it preferentially nominates BGCs that already encode known scaffolds: every one of the five confirmed bioactive products from the five-strain cohort fell within the low-to-mid N band, and a third of the SRE-selected BGCs were near-perfect MIBiG matches that novelBGC would have flagged as rediscovery risks before any experimental commitment. The outcome of this is equally remarkable, as a reservoir of 55 high-N candidates (N ≥ 0.50) was never selected by the SRE filter, including BGCs at the very top of the novelty ladder (e.g. *S. spororaveus* 5.20, N = 0.96; *S. nojiriensis* 1.20, N = 0.88) for which no resistance signature was ever encoded. The lariocidin and indanopyrrole-A case studies (Table 1 rows 6-7) show that the framework also generalises in two complementary directions: the maximal in-genome N-score correctly isolated the productive BGC even in a *Paenibacillus* genome with no MIBiG matches and no known producer family, while a moderately divergent N = 0.59 / RS = 0.31 profile correctly identified the indanopyrrole-A BGC as a worthwhile target for targeted, pattern-based mining without artificially inflating its global rank.

**Figure 4.**
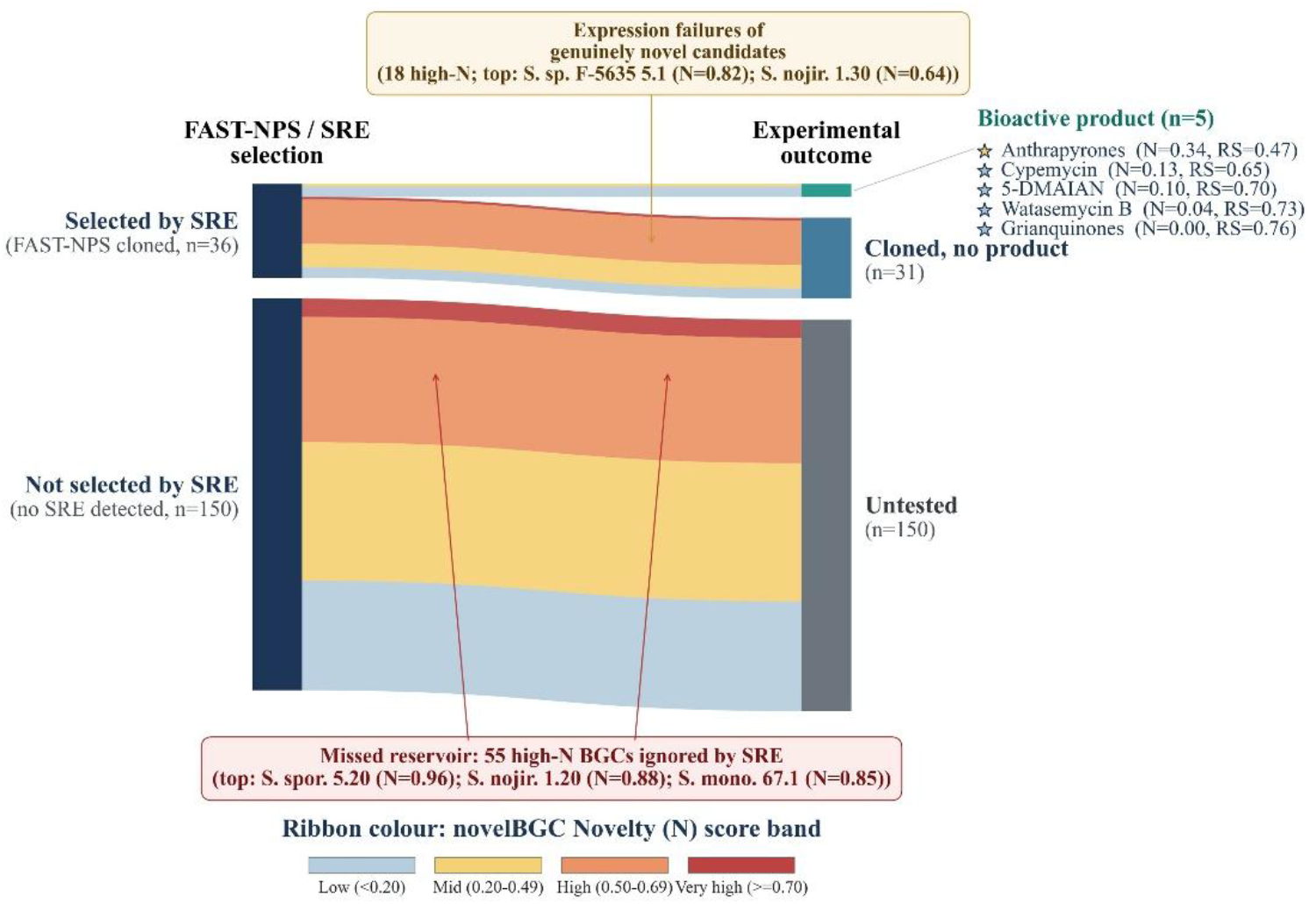
Distribution of FAST-NPS BGCs (n = 186 across 5 *Streptomyces* strains) by SRE-based selection decision and experimental outcome, with ribbons coloured by novelBGC Novelty (N) score band. All five SRE-selected, cloned-and-bioactive BGCs (right-hand panel) fall in the low-to-mid N band, illustrating that the SRE-only filter preferentially selects rediscovery-prone candidates. A reservoir of 55 high-N BGCs (N ≥ 0.50) was never selected by SRE, and a further 18 high-N candidates were cloned but produced no detectable compound. N-band edges: low (<0.20), mid (0.20-0.49), high (0.50-0.69), very high (≥0.70).

Beyond the algorithmic contribution, novelBGC is delivered as a hosted web server, a design choice motivated by a well-documented accessibility gap in modern genome-mining workflows. As Laub and Devraj (2023) note, a substantial fraction of publicly available genomics data is never re-interrogated after its initial publication, not because the underlying biological questions are exhausted, but because wet-lab researchers are seldom formally trained in bioinformatic tooling and therefore assume they lack the experience required to engage with it (Laub et al., 2023). The same authors argue that freely available, predominantly web-based, GUI-driven platforms, requiring no command-line navigation and no local installation, are the most effective route to integrate bench scientists into computational pipelines. novelBGC is built squarely on this principle: it requires no command-line expertise, no local installation of antiSMASH or BiG-SLiCE, and no manual integration of intermediate output formats, presenting instead a single submission page and a single ranked results table. This architectural decision is consequential, because it lowers the practical entry barrier for the very community whose participation matters most for translating predictions into molecules, but who have historically been excluded from genome-mining workflows by infrastructure overhead. To preserve scientific control without re-introducing that overhead, all scoring weights are user-adjustable at the time of submission, so groups with idiosyncratic priorities, for example weighting AMR co-localisation more strongly when chasing antibiotic leads, or weighting GCF distance more strongly when chasing structural novelty, can tune the model without re-implementing it.

Several caveats merit explicit acknowledgement. The N and RS scores are anchored to fixed reference resources (MIBiG and BiG-FAM) and will systematically over-estimate novelty for BGC families that are well described in unindexed literature but absent from those databases; periodic re-indexing against updated MIBiG and BiG-FAM releases is therefore a planned maintenance commitment rather than an optional refresh. The framework does not encode experimental tractability, host expression capacity, or chemical accessibility, and the indanopyrrole-A example (10th by N within its genome) illustrates that a high N-score is a necessary but not sufficient condition for productive prioritisation in scaffold-targeted campaigns. novelBGC also inherits the false-negative rate of antiSMASH for cryptic or fragmentary clusters, particularly RiPPs and unusual hybrid architectures, and the contig-edge penalty, while empirically calibrated for typical short-read assemblies may under-credit genuinely novel clusters that happen to lie at the boundaries of long-read scaffolds. Finally, we caution that the four N-bands used for visualisation in Figure 4 (low, mid, high, very high) are presentational thresholds: the underlying score is continuous, and any operational cut-off should be re-evaluated against the user’s research context rather than treated as a fixed decision boundary.

Future development will focus on integrating transcriptomic and metabolomic evidence to weight predictions by expression, on incorporating learned chemistry-aware embeddings to refine the GCF distance measure, and on supporting user-uploadable resistance-target or scaffold-bait sets so that the global novelty landscape can be intersected with project-specific search criteria. Looking forward, novelBGC reframes BGC prioritisation as a two-dimensional decision solution in which novelty and reference similarity are jointly visualised, transparently weighted, and interactively explorable. Overall, by mathematically synthesising disparate bioinformatics outputs into a continuous two-dimensional landscape, novelBGC not only streamlines current discovery pipelines but also highlights reservoirs of uncharacterized biosynthetic potential. For instance, our retrospective analysis of the *Streptomyces sp*. CNX-435 genome revealed several BGCs with extreme novelty scores (e.g., Region 1.1 at N = 0.95; Region 4.2 at N = 0.93) should not have been bypassed during targeted structural mining and therefore, remain entirely uncharacterized. Such regions represent highly divergent chemical dark matter and by explicitly quantifying this divergence and novelty, novelBGC generates data-driven hypotheses, offering researchers precise coordinates for future experimental exploration.

## Ethics approval and consent to participate

Not applicable

## Data Availability

The authors have used open-source data and tools in this analysis. Information on all used tools is provided in the methodology section.

## Code Availability

Codebase for novelBGC is available from the corresponding author upon reasonable request.

## Authors’ contributions

GShu and GSha conceptualized the idea. GSha supervised the project. GShu performed the analysis and wrote the first draft. BM helped in analysis. All authors edited, finalized, and approved the final version of the manuscript.

